# Tree and shrub richness modify subtropical tree productivity by modulating the diversity and community composition of soil bacteria and archaea

**DOI:** 10.1101/2022.07.30.502115

**Authors:** Siqi Tao, G. F. (Ciska) Veen, Tianhe Yu, Naili Zhang, Laiye Qu

## Abstract

**Background:** Declines in plant biodiversity often have negative consequences for plant community productivity, and it becomes increasingly acknowledged that this may be driven by shifts in soil microbial community composition. These relationships have been well-established in grasslands, and few studies also indicate that fungi play a role in driving tree diversity-productivity relationships in forests. However, the role of bacteria and archaea, which are also highly abundant in forest soils and perform pivotal ecosystem functions, has been largely overlooked. Here, we investigated how tree and shrub richness affects stand-level tree productivity via modulating bacterial and archaeal community diversity and composition. We used a landscape-scale, subtropical tree biodiversity experiment (BEF-China) where tree (1, 2 or 4 species) and shrub richness (0, 2, 4, 8 species) were modified.

**Results:** Our results showed that tree and shrub species richness affected bacterial diversity, community composition, and co-occurrence networks, but such effects were marginal for archaea. Both tree and shrub species richness increased stand-level tree productivity by modulating soil microbial community composition, with the effects being mediated via increases in soil C:N ratios.

**Conclusions:** Our findings imply the importance of bacterial and archaeal communities in driving the relationship between plant diversity and productivity in subtropical forests and highlight that we may require better a understanding of prokaryotic communities in forest soils.

## Background

Anthropogenic activities have resulted in the loss of biodiversity worldwide [1–2] and in altered ecosystem functioning [3] and the services that ecosystems provide to humanity [4]. This has fostered a large research field that aims at understanding the relationship between biodiversity and functioning [5]. Much of the work originates from grassland systems [6–9], where it has been found that plant species diversity generally increases plant community productivity and that this relationship is driven by shifts in the soil community [10]. Although recent studies found similar biodiversity-ecosystem functioning (BEF) relationships in forests [11–14], it is still poorly understood how changes in soil biodiversity contribute to increased productivity in diverse tree stands. Moreover, the presence of shrubs in forests can interfere with the diversity effects of trees. In general, shrubs in the understory may reduce tree productivity [15–16], but these effects may be reduced at higher levels of shrub richness [16]. Therefore, to fully understand BEF relationships in forests it will be of importance to test how tree and shrub diversity in forest ecosystems drive soil community composition and tree productivity.

Soil biodiversity may underlie positive effects of biodiversity on productivity, because a higher plant diversity may result in a higher variety of plant-derived resources, which in turn will enhance multi-trophic diversity [17] and thus ecosystem functioning [18–19]. Evidence from long-term diversity experiments in grassland support the idea [20–21] that plant diversity drives the structure and functioning of soil microbial communities through the bottom-up (resource control) effects [7, 20]. In forests, fungi are known to form symbiotic relationships with plants in forest soils [22–24] and fungal community composition is closely linked to tree diversity [25–28]. However, bacteria and archaea are also abundant in forest soils [29] and are known to play key roles in carbon fluxes and nutrient cycling and decomposition [30–37], but to what extent shifts in bacterial and archaeal diversity, community composition and community complexity (e.g., network structure, connectedness etc.) underlie BEF relationships in forests has not been tested yet. In addition, there is limited information on whether the mycorrhizal types of the focal tree species shape soil prokaryote communities in the context of changing tree and shrub species richness levels, although there has been extensive research demonstrating that it has a significant effect on fungal communities [28, 38–39].

As plant diversity increases, plant-plant interactions develop, and so does the complexity of the associated microbe-microbe interactions. Given that non-random community assembly may be a general characteristic for microorganisms [40], correlation-based network of co-occurring microorganisms based on strong and significant correlations (non-parametric Spearman’s) [41] was widely used to reveal microbial co-occurrences and the connectivity among community members [42–47]. And a pioneer research in experimental grassland ecosystems observed that microbial network complexity positively influences multiple ecosystem functions [48]. It would be of great interest to examine whether changes in plant diversity could influence the microbiome complexity, such as diversity and interconnectedness among co-occurring microbes, and whether it could have impact on the way in which microbe communities influence ecosystem function.

To investigate how soil prokaryotic community composition, diversity and co-occurrence networks respond to tree and shrub species richness and how this in turn affects tree productivity, we conducted an experiment in BEF-China platform: a subtropical forest in southeast China where tree and shrub diversity are experimentally varied [49]. We used Illumina amplicon sequencing of small subunit ribosomal RNA markers to determine the communities of bacteria and archaea in bulk soils under the canopy of focal trees. Along the three tree species richness levels (1, 2, 4) with four shrub species richness levels (0, 2, 4, 8), we investigated the relationships between plant diversity with bacterial and archaeal diversity, composition, and co-occurrence relationships, furthermore, how microbes respond to changes in aboveground plant diversity and thus regulate stand-level tree productivity. We hypothesized that: (H1) tree and shrub species richness would interactively affect bacterial and archaeal diversity and composition, microbial *alpha* diversity increases with the increasing tree species diversity but decreases with the increases of shrub species diversity; (H2) the microbial network complexity would increase with tree and shrub species diversity; (H3) tree and shrub species diversity would promote the stand-level tree productivity through modulating the composition and diversity bacterial and archaeal communities.

## Methods

### Study area

The BEF-China platform (https://bef-china.com/) has been set up to investigate the relationship between subtropical plant diversity and ecosystem functioning in Xingangshan, Jiangxi Province in southeast China (29°08’–29°11’ N, 117°90’–117°93’ E) [49]. It took for two years from 2009 to 2010 to finish the establishment of the main experimental sites of BEF-China platform, where it belongs to the subtropical climate zones. The mean annual temperature is 16.7 °C, with the coldest temperature 0.4 °C occurred in January, and the warmest 34.2 °C in July [50], while mean annual precipitation is 1821 mm. The vegetation in natural ecosystems surrounding the BEF-China platform is an evergreen and deciduous broad-leaves mixed forest [51]. The soils belong to Regosols, Cambisols, Acrisols, Gleysols and Anthrosols [52].

### Experimental setup and sampling

The design of BEF-China main experiment was described by [49]. Specifically, two experimental sites, A (18.4 ha) and B (20 ha), were respectively set up on the field after clear-cutting *Pinus massoniana* and *Cunninghamia lanceolata* plantation. There were 32 4 mu super-plots that were divided into four plots with a size of 25.8 × 25.8 m (1 mu of Chinese area unit) in both sites. Within each plot, there were 400 trees randomly planted in 20 × 20 grids, with a 1.29-m interval between tree individuals along the cardinal compass directions. A species pool containing 40 broadleaved tree species and 18 shrub tree species was first established, to minimize the confounding effects of a particular species combination on diversity effects [49]. Based on the species pool, tree and shrub species were randomly selected to build a crossed tree and shrub species richness gradient. The super-plots represented five tree species richness levels: one- (16 super-plots), two- (8 super-plots), four- (4 super-plots), eight- (2 super-plots) and sixteen- (1 super-plot) and twenty-four species richness (1 super-plot). There were 32 4 mu super-plots in total, with 128 1 mu plots. Within each 4 mu super-plot, there were four 1 mu plots where shrubs were planted, with 0, 2, 4 or 8 shrub species richness randomly assigned in these plots.

In this study, 56 plots with three tree species richness levels (1, 2, 4) coupled with four shrub species richness levels (0, 2, 4, 8) in both site A and B were chosen in October 2018 (Figure 1a, b). Four focal tree individual per species in each plot were randomly chosen, and a composite soil sample was collected per individual (Additional file Figure S1). Specifically, four soil cores at the 0-10 cm depth were collected from different directions within 1/2 of the canopy projection area of each tree individual, and well mixed to avoid spatio-temporal autocorrelation. As such, a total of 192 samples were yielded in each site (Additional file Table S1). These soil samples were further divided into two parts: 1) air-dried for soil physicochemical properties measurement; 2) stored at -80 °C for DNA extraction and subsequent microbiome analyses.

**Figure 1.**
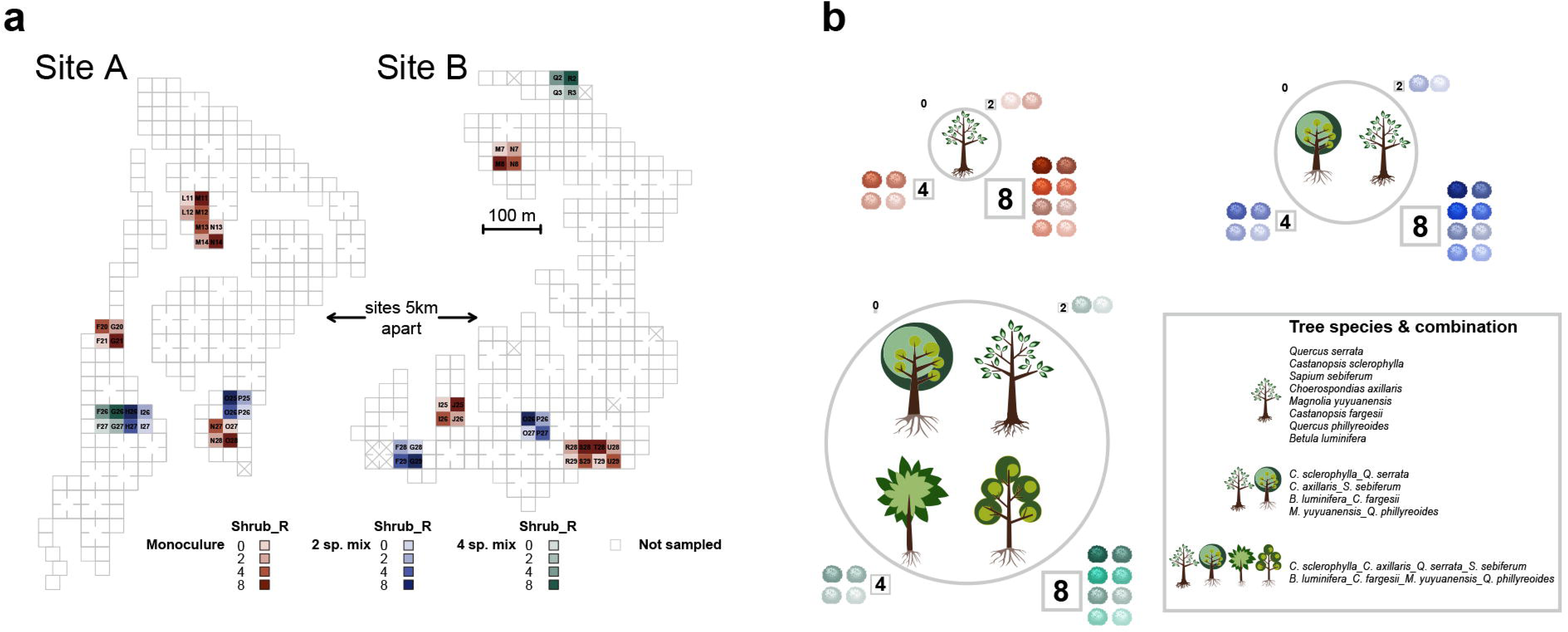
Sampling and experimental design. a. Plots with tree species richness gradients (1, 2, 4) and shrub species richness gradients (0, 2, 4, 8) selected from BEF-China platform (site A, site B). b. The tree species and their combinations used in this study.

### Topographic and soil physicochemical properties

A digital elevation model was used to estimate mean plot aspect and inclination as described by [53–54]. Two components, i.e., a north–south and an east–west slope aspect, were calculated based on the mean plot aspect. Because there are also plots in flat areas and on small slopes that cannot be classified as a particular aspect, hence two identified components d.SLOPEnew and d.GRA_NS used in this study were estimated according to the following equations:

d.SLOPEnew = SLOPE*pi/180 d.GRA_NS = tan(d.SLOPEnew)*NS

Fresh soil samples were sieved with a 2-mm sieve to measure soil properties. Soils were dried at 105 °C for 48 h for determining soil water content. Soil solutions with 1: 2.5 soil to water ratio were used to measure soil pH with a glass electrode (Thermo Orion T20, USA). Air-dried soils were used to estimate soil organic C and total N with the CHNOS Elemental Analyzer (vario EL III, CHNOS Elemental Analyzer; Elementar Analysensysteme GmbH, Langenselbold, Germany). Soil P and other chemicals, i.e., calcium (Ca), potassium (K), magnesium (Mg), and ferrum (Fe) were measured with inductively coupled plasma emission spectrometry (ICAP 6300 ICP-OES Spectrometer; Thermo Scientific, Waltham, MA, USA). Soil inorganic N, i.e., nitrate and ammoniacal N were measured using Continuous Flow Analyzer (SAN Plus, Skalar, Erkelenz, Germany).

### Tree stand volume and increment

Tree stand volume and increment data were retrieved from [55] to estimate the stand-level tree productivity. Briefly, individual tree volume proxies were calculated as H × π (BR)^2^ in which H is height and BR is basal radius at the ground, and then transformed to more accurate tree volume estimates by multiplying the proxies with a size-dependent correction factor according to [16]. The stand-level tree volume was calculated by aggregating the volumes of the surviving trees in the central 36 planting positions per plot and stand volume increment was calculated as the absolute differences in stand volume between two consecutive years. In our study, we used the tree stand volume data in 2018 and volume increment data between 2018 and 2019.

### Soil microbial biomass

Microbial biomass was measured by the chloroform fumigation extraction method [56]. A pair of fresh soils per sample with 5 g weight of each were separately added into beakers, and then one of them was placed into a vacuum drier with 50 ml alcohol-free CHCl_3_ to fumigate for 24 h, while the other was assigned as the control without fumigation. The paired fumigated and un-fumigated soils were both incubated at 25 °C for 24 h in the dark. The paired soils were extracted using 50 ml 0.5 M K_2_SO_4_ (1:2.5 w/v), and then C and N concentration in soil solutions were measured with TOC analyzer (Liqui TOC II; Elementar Analysensysteme GmbH, Hanau, Germany). The formula calculating microbial biomass C (MBC) and microbial biomass N (MBN) is as followed: B_c(n)_ == F_c(n)_ /k_c(n)_ . Here, F_c(n)_ referes to difference value between amount of C or N extracted from fumigated and unfumigated soil. k_c(n)_ refers to the calibration coefficient of microbial biomass, where k_c_ is 0.38 for MBC [57] and k_n_ is 0.54 for MBN [56].

### DNA extraction, PCR amplifications and sequencing

Soil samples packed with dry ice were transferred and stored at -80°C in laboratory until DNA extraction. The extraction of microbial genomic DNA was conducted using the PowerSoil DNA Isolation Kit (Mobio, Laboratories, Inc., Carlsbad, CA, USA) according to the manufacturer’s protocols. The concentration of DNA extracts was determined using the NanoDrop 2000 UV-vis spectrophotometer (Thermo Scientific, Wilmington, USA), and the quality of DNA extracts were examined using 1% agarose gel electrophoresis. The primer pairs 338F (5’-ACTCCTACGGGAGGCA GCAG-3’) and 806R (5’-GGACTACHVGGGTWTCTAAT-3’) [58–59], and the primer pairs 524F10extF (5’-TGYCAGCCGCCGCGGTAA-3’) and Arch958RmodR (5’-YCCGGCG TTGAVTCCAATT-3’) [60] were used to amplify the hypervariable region V3-V4 of the bacterial 16S rRNA gene and V4-V5 of archaeal 16S rRNA gene, respectively.

The PCR amplification of 16S rRNA gene was performed as follows: initial denaturation at 95 °C for 3 min, followed by 27 cycles of denaturing at 95 °C for 30 s, annealing at 55 °C for 30 s and extension at 72 °C for 45 s, and single extension at 72 °C for 10 min, and end at 10 °C. The PCR mixtures contain 5 × *TransStart* FastPfu buffer 4 μL, 2.5 mM dNTPs 2 μL, forward primer (5 μM) 0.8 μL, reverse primer (5 μM) 0.8 μL, *TransStart* FastPfu DNA Polymerase 0.4 μL, template DNA 10 ng, and finally ddH_2_O up to 20 μL. The PCR reactions were performed in triplicate. The PCR products were extracted from 2% agarose gel and purified using the AxyPrep DNA Gel Extraction Kit (Axygen Biosciences, Union City, CA, USA) according to manufacturer’s instructions and quantified using Quantus™ Fluorometer (Promega, USA). The qualified PCR products were mixed, and paired-end sequenced on an Illumina MiSeq PE300 platform (Illumina, San Diego, USA) according to the standard protocols by Majorbio Bio-Pharm Technology Co. Ltd. (Shanghai, China).

### Bioinformatics analysis

All paired rRNA amplicon sequencing raw reads were processed via the Quantitative Insights into Microbial Ecology 2 (QIIME2) version 2020-2 [61]. Briefly, raw sequence data were imported into QIIME2 manually using the “qiime tools import” command. The quality trimming, denoising, merging and chimera detection and non-singleton amplicon sequence variants (ASVs) grouping were done using the plugin “qiime dada2 denoise-paired” in DADA2 [62] as implemented in QIIME2 v2020-2. The “-p-trim-left-f” and “-p-trim-left-r” parameters were set at 0 and the “-p-trunc-len-f” and “-p-trunc-len-r” parameters were set at 294 for bacteria and 298 for archaea, respectively, after reviewing the “Interactive Quality Plot tab” in the “demux.qzv” file. After the quality filtering steps, the ASV abundance tables were rarefied at 4337 for bacteria and 2083 for archaea, according to the “Interactive Sample Detail” in the “table.qzv” file, respectively to ensure even sampling depth. The *alpha* diversity analyses were conducted from the rarefied ASV abundance tables through the core-metrics-phylogenetic method in the q2-diversity plugin. The bacteria AVSs were taxonomically classified using the qiime2 v2020-2 plugin “qiime feature-classifier classify-sklearn” with the pre-trained Naïve Bayes Greengenes classifier trimmed to the V3-V4 region of the 16S rDNA gene. The archaea ASVs were analyzed by RDP Classifier [63] against the SILVA Small Subunit rRNA Release v11.5 using a confidence threshold of 0.7. Furthermore, the taxa that were not present in at least 5% of total samples were removed from the matrices for both bacteria and archaea to reduce the noise. The bacterial and archaeal ASVs were functionally annotated by FAPROTAX [64] and assigned to putative functional groups, i.e., microbial groups associated with carbon cycle, nitrogen cycle or sulphur cycle.

### Statistical analysis

All the statistical analyses and data visualization were performed in R (Version 3.6.3). *Beta* diversity was assessed by computing unweighted UniFrac distance matrices using the vegan package [65]. The significance of different factors on community dissimilarity was tested with PERMANOVA by permutations of 999 in using the ‘adonis’ function of the vegan package [65] based on unweighted UniFrac distances. Statistically significant differential abundant taxa among different species richness were determined with the DESeq2 package [66] using pair-wise comparisons in a negative binomial generalized linear model at an FDR-adjusted *p* value of 0.05. Bipartite networks were generated using Cytoscape [67] following the method in [68]. The network analysis was performed by igraph package [69] and visualized in Gephi [70] to explore the co-occurrence of features from a holistic perspective. The correlation between the community diversity distance matrix and topographic/soil physicochemical properties matrix was assessed from Mantel tests and visualized by MatCorPlot package [71]. To tease apart the effects of tree and shrub species richness on bacterial or archaeal microbiome and the consequences on stand-level tree productivity, Structural Equation Modelling (SEM) were performed. The SEM models were built based on the conceptual model shown in Additional file Figure S2, using the “sem” function in lavaan package [72]. The path coefficient represents the direction and strength of the direct effect between two variables. The goodness of fit was estimated using three indices: (i) the root mean square error of approximation (RMSEA<0.05), (ii) the comparative fit index (CFI > 0.95), and (iii) the standardized root mean squared residuals (SRMR< 0.08) [73].

## Results

### 1. Soil bacterial and archaeal *alpha* diversity

For the soil bacterial community, *alpha* diversity (expressed as Chao1) reduced from monocultures to 2-tree species mixtures (*p* < 0.001) and 4-tree species mixtures (*p* < 0.001) and was affected by an interaction between shrub and tree species richness (Figure 2a). The interaction indicated that bacterial *alpha* diversity increased with increasing shrub species richness for tree monocultures but decreased in 4-tree species mixtures. When tree species richness is 2, *alpha* diversity increased generally with increasing shrub richness, except for a significant decrease in shrub richness at 4. For archaea, Chao1 increased from monocultures to 2-tree species mixtures (*p* < 0.05) and 4-tree species mixtures (*p* < 0.05) (Figure 2a). Moreover, shrub species richness enhanced archaeal *alpha* diversity, especially in the context of 2- and 4-species tree mixtures (Figure 2a). When considering the mycorrhizal types of the focal tree species, we found the bacterial diversity was higher for ectomycorrhizal (EcM) than for arbuscular fungi-colonized trees (AM) (*p* < 0.05). The bacterial diversity decreased from monocultures to polycultures for both EcM and AM trees (*p* < 0.001) (Additional file Figure S3a). However, no significant differences of archaeal diversity were found between EcM and AM tree species, only the archaeal diversity of EcM tree species increased with increasing tree species richness (*p* < 0.01), but not for AM tree species (Additional file Figure S3b).

**Figure 2.**
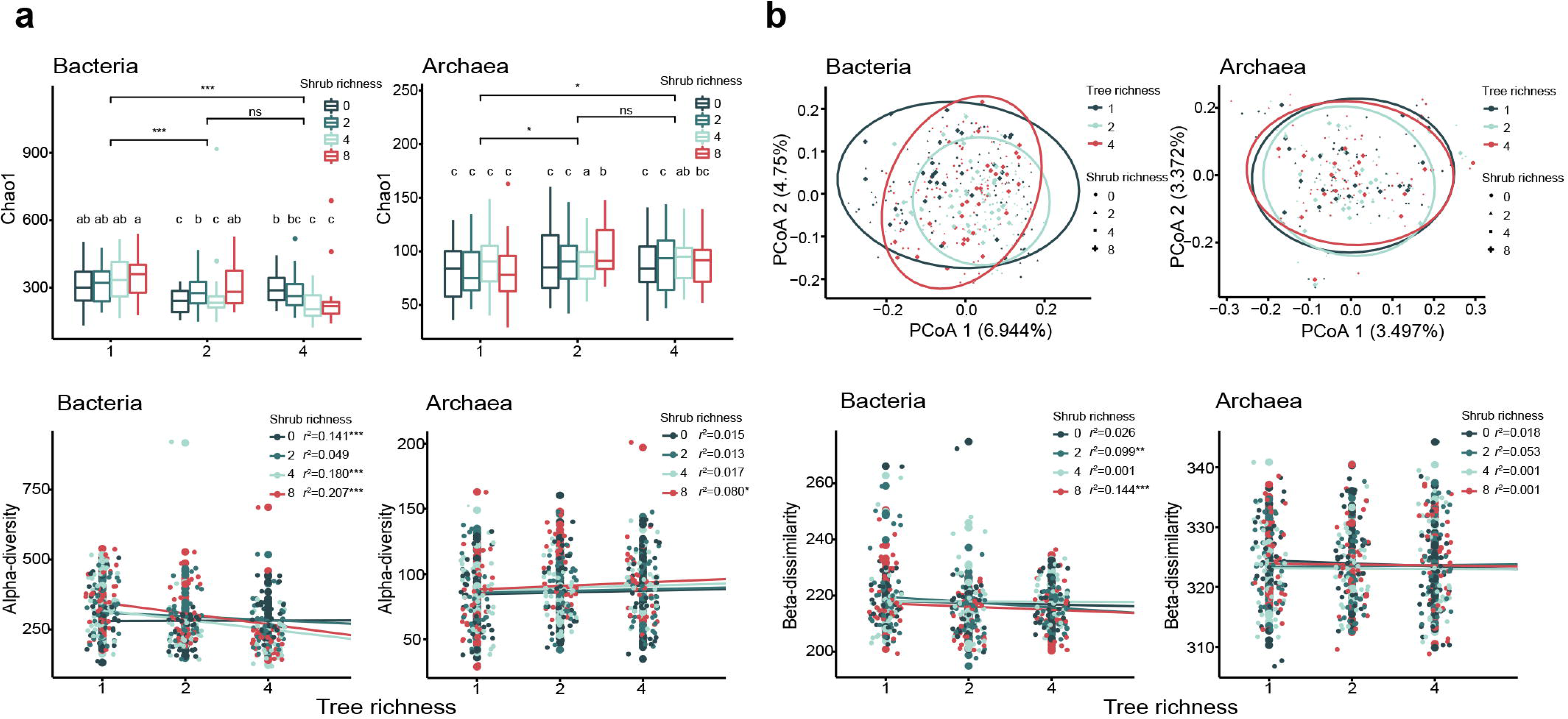
Soil bacterial and archaeal *alpha* diversity and community structure. a. tree richness and shrub richness effects on soil microbial *alpha* diversity; b. tree richness and shrub richness effects on soil microbial *beta* dissimilarity. The asterisks showed the p-value significance level, *p < 0.05, **p < 0.01, ***p < 0.001, ****p< 0.0001 and ns showed no significance.

We found that the ratio of MBC to soil organic C (MBC/C_org_) and the ratio of MBN to soil organic N (MBN/N_org_) were positively correlated to archaeal *alpha* diversity. Soil organic C significantly negatively related to both bacterial and archaeal *alpha* diversity (Table 1).

**Table 1.** Pearson correlation of bacterial and archaeal diversity with environmental variables. SM, MBC, MBN, C_org_ and N_org_ respectivly refers to soil moisture, microbial biomass C, microbial biomass N, soil organic C and N.

|  | Bacterial |  | Archaeal |  | Bacterial |  | Archaeal |  |
| --- | --- | --- | --- | --- | --- | --- | --- | --- |
|  | abundance |  | abundance |  | Chao1 index |  | Chao1 index |  |
|  | r | p | r | p | r | p | r | p |
| pH | -0.084 | 0.099 | -0.017 | 0.742 | -0.060 | 0.241 | -0.033 | 0.523 |
| SM | -0.006 | 0.900 | -0.037 | 0.465 | -0.010 | 0.850 | -0.057 | 0.267 |
| NH<sub>4</sub><sup>+</sup> | -0.040 | 0.434 | 0.048 | 0.351 | -0.015 | 0.762 | 0.048 | 0.344 |
| NO<sub>3</sub><sup>-</sup> | 0.067 | 0.193 | 0.113 | **0.026** | 0.060 | 0.237 | -0.085 | 0.096 |
| N | -0.010 | 0.851 | 0.084 | 0.099 | 0.009 | 0.854 | 0.010 | 0.847 |
| MBC | 0.047 | 0.360 | 0.104 | **0.042** | 0.072 | 0.161 | 0.030 | 0.557 |
| MBN | 0.022 | 0.668 | -0.012 | 0.811 | 0.003 | 0.947 | -0.009 | 0.856 |
| MBC/MBN | 0.004 | 0.934 | 0.060 | 0.243 | 0.026 | 0.607 | 0.006 | 0.901 |
| MBC/C<sub>org</sub> | 0.081 | 0.115 | 0.176 | **0.001** | 0.115 | **0.025** | 0.121 | **0.017** |
| MBN/N<sub>org</sub> | 0.047 | 0.358 | 0.113 | **0.027** | 0.047 | 0.360 | 0.107 | **0.037** |
| P | -0.052 | 0.309 | -0.051 | 0.316 | -0.065 | 0.203 | -0.034 | 0.513 |
| C | -0.125 | **0.014** | -0.165 | **0.001** | -0.142 | **0.005** | -0.189 | **0.000** |
| C/N | -0.002 | 0.976 | 0.137 | **0.007** | 0.025 | 0.631 | 0.059 | 0.252 |

### 2. Bacterial and archaeal community composition

We found that bacterial community composition differed between levels of tree species richness and shrub species richness, but the effects of tree species richness on bacterial community composition were stronger than effects of shrub species richness (Figure 2b; Table 2). In contrast, soil archaeal communities were influenced by shrub species richness, but generally not by tree species richness (Figure 2b; Table 2). In addition, bacterial community composition was influenced by the interaction between tree and shrub species richness (Table 2).

**Table 2.**
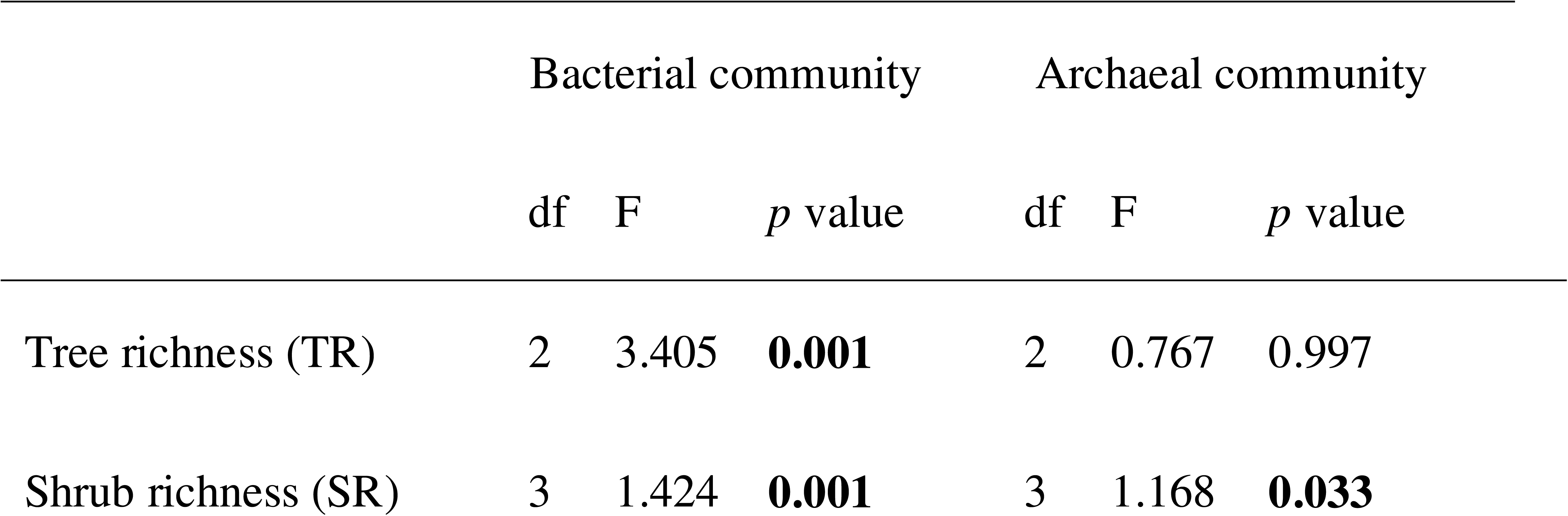
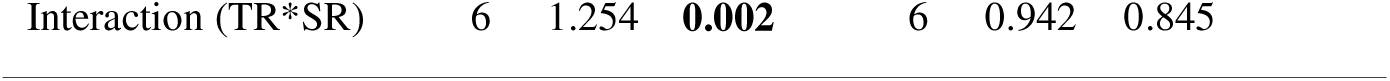
The effects of tree and shrub richness on the compositional variances of soil bacterial and archaeal communities based on PERMANOVA with 999 permutations.

Soil bacterial community composition varied between EcM and AM trees (PERMANOVA test, F=1.68 *p* < 0.010) (Additional file Figure S4a). In addition, tree species richness had a significant effect on bacterial community structure under both EcM and AM trees (Additional file Table S2). For archaeal community composition differences between tree mycorrhizal types were less pronounced (Additional file Figure S4b) and it was not affected by differences in tree and shrub species richness under the canopy of EcM or AM trees (Additional file Table S2).

We found that soil moisture (SM), pH, the soil C/N and two topographical factors d.SLOPEnew and d.GRA_NS were positively associated with bacterial community composition (*p* < 0.05) (Additional file Figure S5). However, there was no significant correlation between these factors and soil archaeal community composition (Additional file Figure S5).

### 3. Bacterial and archaeal taxonomic and functional groups

For all 384 soil samples, we obtained a total of 70,836 ASVs for bacterial and 13,552 ASVs for archaeal communities. The dominant bacterial phyla across all samples were Acidobacteria (36.44% of the total bacterial sequences), Proteobacteria (27.99%) and Chloroflexi (6.37%) (Figure 3a). As for the taxonomic abundance of the soil archaeal communities, the phyla Thaumarchaeota (56.97% of the total archaeal sequences), Euyarchaeota (29.00%) and Crenarchaeota (12.78%) dominated the archaeal communities (Figure 3a).

**Figure 3.**
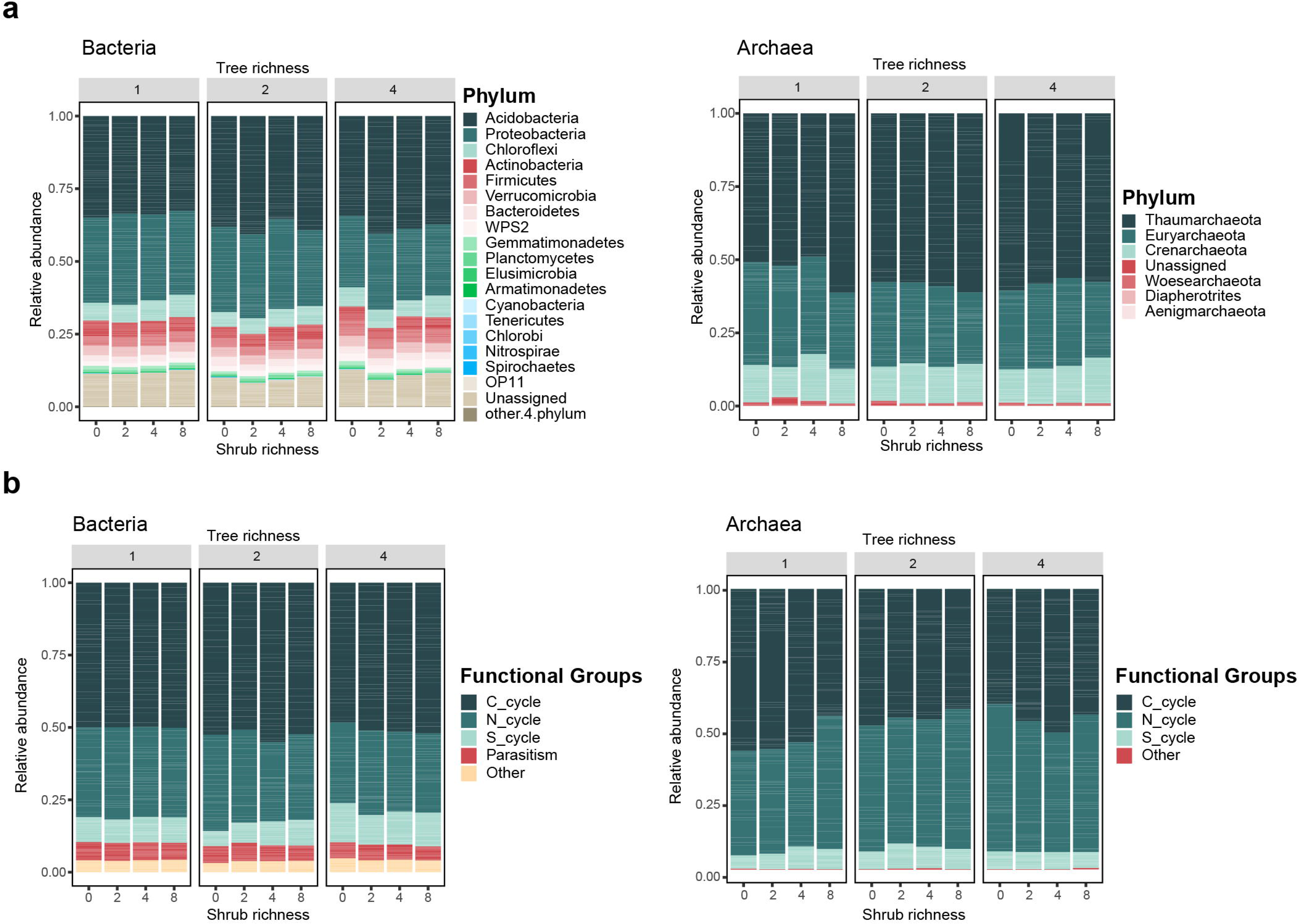
Taxonomic and functional classifications of soil bacterial and archaeal communities. a. the relative of phylum-level taxa dominated across tree species richness and shrub species richness levels; b. functional assignments with relative abundance of each functional groups in bacterial and archaeal community, including carbon cycling (C_cycle), nitrogen cycling (N_cycle), sulfur cycling (S_cycle), parasitism and others under the combined effects of tree species richness and shrub species richness.

We found that tree species richness affected the relative abundance of certain bacterial taxonomic groups (Additional file Table S3). For example, the relative abundances of Chloroflexi increased with increasing tree species richness, while the relative abundances of Acidobacteria and Firmicutes were lower in 4-species mixtures compared to monocultures and 2-species mixtures. The abundance of the phylum Proteobacteria was lower in 2-species mixtures than monocultures (Additional file Table S3). In addition, the relative abundance of bacterial phyla was more likely to change with increasing shrub species richness, as tree species richness increased (Additional file Table S3). When tree species richness was at level 1, the relative abundance of bacterial phyla did not differ between shrub monocultures and other shrub diversity levels. However, the relative abundance of Acidobacteria decreased significantly from shrub richness level 2 to 4 when tree species richness level increases to 2, and increased significantly from shrub richness level 2 to 4 (or 8) when tree species richness increased to 4 (Additional file Table S3). For archaea we found little effects of tree and shrub species richness on relative abundances (Additional file Table S3). Furthermore, we did not find significant differences in the taxonomic composition of bacterial or archaeal communities in soils collected under EcM and AM trees (Additional file Figure S6).

Our results showed that a large proportion of bacterial and archaeal ASVs were assigned to C-cycle and N-cycle groups (Figure 3b). The relative abundance of bacteria was significantly lower in the C-cycle and S-cycle groups but higher in the N-cycle group in 4-tree species mixtures compared to monocultures and the relative abundance of archaeal C-cycle group was significantly higher in monocultures than 2-tree species mixtures (Additional file Table S4).

### 4. Bacterial and archaeal network complexity

Tree species richness decreased network complexity for bacteria (Figure 4a), indicated by a decline in the average degree, network density, modularization, the number of nodes and the number of edges. Network complexity did not change for archaea (Figure 4b) and interactive effects of tree and shrub species richness on soil bacterial and archaeal co-occurrence networks were limited (Additional files Figure S7-S8).

**Figure 4.**
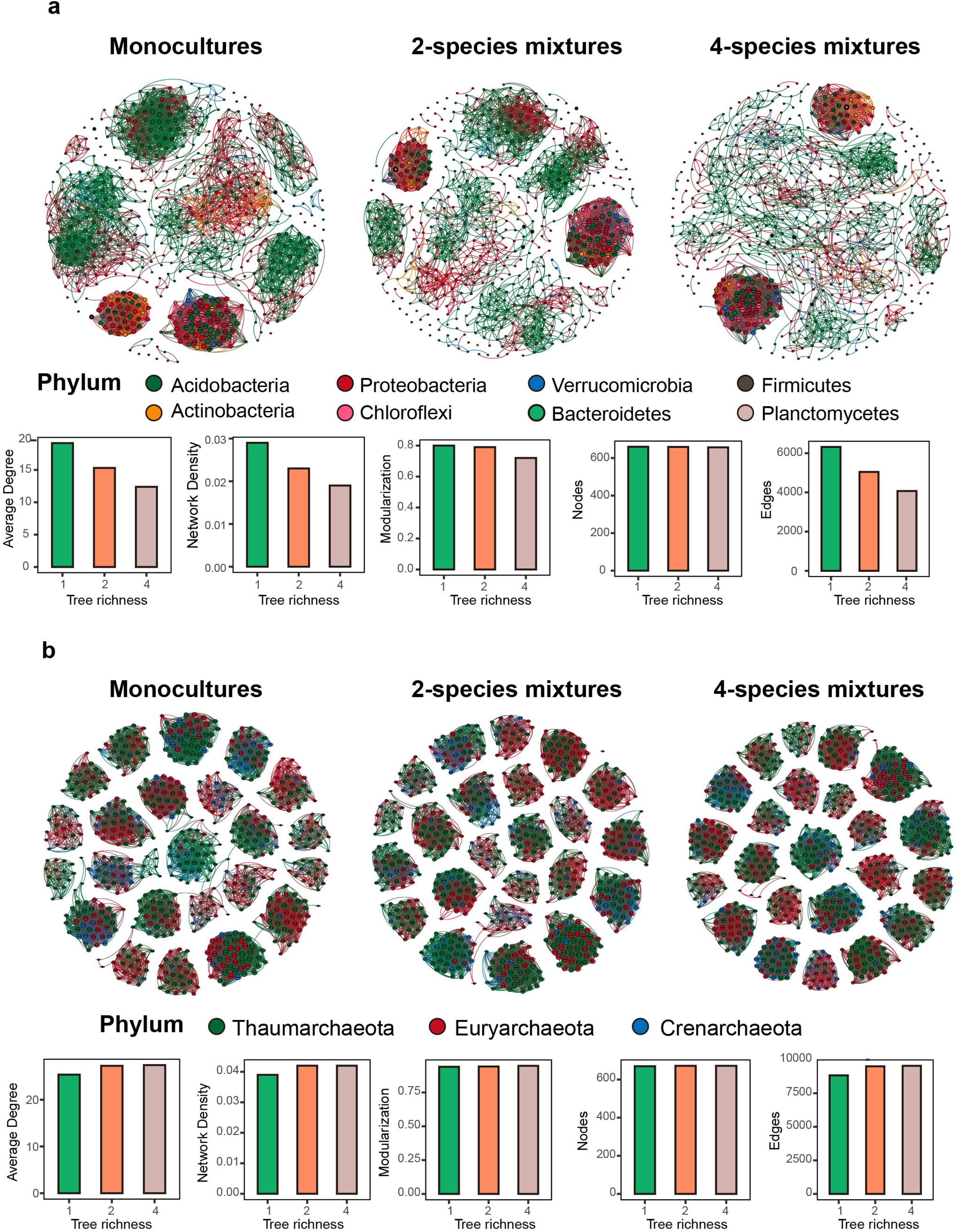
The co-occurrence networks of bacterial communities (a) and archaeal communities (b) in three tree species richness levels, monocultures, two-species mixtures and four species-mixtures, respectively. The nodes in the networks are colored according to the taxonomic assignments at phylum level and the size of each node is proportional to the relative abundance.

### 5. The role of soil bacteria and archaea modifying BEF relationship

Using SEM, we found that both tree and shrub species richness had a positive effect on soil C/N and stand-level tree productivity (Figure 5). Soil C/N was positively correlated with MBC/C_org_, and consequently increased tree productivity (Figure 5).

**Figure 5.**
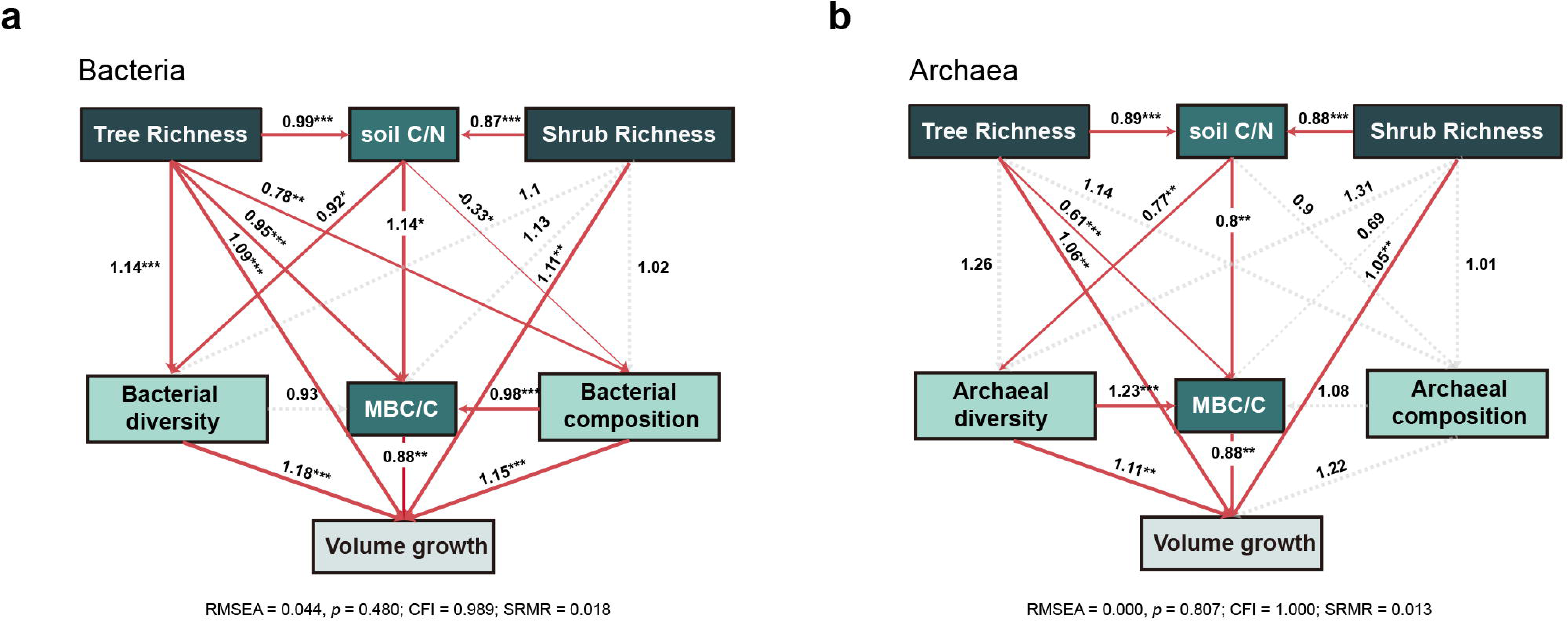
Structural equation models demonstrating the direct and indirect effects of aboveground plant richness on soil nutrient contents, microbial biomass and community-level tree productivity, red arrows indicate significant and positive relationships (*p* < 0.05) and dashed arrows indicate connections with insignificant relationship (*p* > 0.05).

Tree species richness exhibited a significantly positive effect on MBC/C_org_ as well (Figure 5). Most interestingly, we found that tree species richness positively linked to bacterial diversity, and modulate bacterial community composition, which then contributed to the increase in stand-level tree productivity (Figure 5a). Impacts of tree species richness on bacterial community composition were modulated via altered soil C/N (Figure 5a). Here, we note that the bacterial composition rather than diversity was a direct positive driver on MBC/C_org_, thereby contributing to an increase in stand-level tree productivity (Figure 5a). Neither tree nor shrub species richness directly altered the diversity and composition of archaeal community (Figure 5b). However, we found that tree species richness influenced archaeal diversity via regulating soil C/N (Figure 5b). Archaeal diversity was positively associated with MBC/C_org_ ratio, which then increased stand-level tree productivity (Figure 5b).

## Discussion

In our study we explored how tree and shrub species richness affected the diversity, complexity, and composition of bacterial and archaeal communities in a large subtropical tree biodiversity experiment. In addition to earlier work on fungal communities [74–75], we now show for the first time that tree species richness drives shift in bacterial and archaeal *alpha* diversity and bacterial community composition (H1). In addition, we found significant interactions between tree and shrub species richness levels, indicating that the shrub species richness effect on bacterial *alpha* diversity was dependent on tree species richness (H1). The complexity of the bacterial networks was found to decrease significantly with increasing tree species richness but was not significantly influenced by changes in shrub species richness (H2). The complexity of archaeal co-occurrence network was not correlated with either tree or shrub species richness (H2). Contrary to the view that the presence of shrub competition in forests may reduce tree productivity [16], we found that both tree and shrub species richness contributed to tree productivity and highlighted soil bacterial and archaeal communities as vital linkages between plant richness and stand-level tree productivity in the context of plant-created soil chemical properties (H3). In summary, our study provided novel insights that diversity and composition of prokaryotic communities are responsive to tree species richness and appear to play a role in driving tree productivity, hence the inclusion of them in forest soil community analyses is therefore important for understanding the functioning of these ecosystems.

### Tree-shrub species richness affected the bacterial and archaeal diversity and community composition under the canopy of focal tree species

In contrast with our first hypothesis that soil microbial *alpha* diversity increases with the increasing tree species diversity, we found that plant community richness had a negative effect on soil bacterial diversity under the canopy of focal trees, indicating that the most diverse bacterial communities in our study occurred in monocultures and that diversity decreased with increasing community-level tree richness. This is in contrast with earlier work in grassland [10] and on fungal communities in the BEF-China experiment [26]. Unlike these earlier studies, we have collected soil samples underneath individual trees rather than at the community-level, and it is therefore reasonable to suggest that the decline in soil bacterial diversity may point to a ‘dilution effect’ [23, 76]. From this perspective, the tree species richness gradient from 1, 2 to 4 resulted in smaller densities of conspecific tree species in the focal tree species, so that some focal tree-specific bacteria may be restricted. In addition to the sampling strategy, we also speculated that the soils in tree monocultures with low-diversity resources may amplify bacterial competitive pressures, resulting in highly antagonistic bacterial communities, while higher diverse plant communities that provided diverse resources to the soil may reduce microbial competitive pressure and generate less diverse bacterial communities [77–79].

In line with our first hypothesis, we found that tree and shrub species richness resulted in shifts in bacterial community composition. In addition, we found that bacterial community composition became more similar with increasing tree species richness, which is in line with earlier findings from the BEF-China study that fungal community composition was more similar in multi-tree species mixtures [28]. The community composition and diversity can be pronouncedly changed by modulating the soil chemistry resources, which can promote or inhibit the relative abundance of specific microbial taxa [80].

As part of our hypothesis (H1), we postulated that the *alpha* diversity of soil archaea increases with increasing tree species richness but decreases with increasing shrub species richness. Our results are partly consistent with this hypothesis that *alpha* diversity of the soil archaea consistently increased with increasing tree species richness, likely due to changes in the abundance of ammonia-oxidizing archaea resulting from increasing tree species richness [81–82]. However, we found that the effect of shrub species richness on archaeal diversity was rather weak. One possible explanation is that the changes in nitrogen content brought about by the changes in shrub species richness were not sufficient to cause significant differences in archaeal diversity. Furthermore, neither tree species richness nor shrub species richness showed significant effect on soil archaeal compositional variation, unlike bacteria, which may be related to their large differences in environmental adaptations, cellular structure, or cellular metabolisms [83–84]. Despite the key role archaea play in soil biogeochemical cycles, studies on how their abundance is influence by plant diversity remain extremely sparse [85].

Notably, our findings also underline the need to consider the tree mycorrhizal types as important factor in studying ‘tree-shrub diversity-soil prokaryotic community’ relationships. We found that both bacterial and archaeal *alpha* diversity showed significant differences between mycorrhizal types and the mycorrhizal type of the focal tree species influenced the microbial response pattern to tree species diversity. This is mainly because different mycorrhizal types-associated fungi differ in their strategies of resource acquisition, nutrient allocation, and plant-soil feedback, which could affect their recruitment of different microbes in the mycorrhizosphere [86]. In contrast to our results, a previous study examining the same field experiment showed no significant difference in soil bacterial *alpha* diversity between EcM and AM trees [28]. This contrasting result may be because the study selected two adjacent EcM and AM trees as a target sampling unit, making the difference in soil nutrient resources not significant enough to affect bacterial diversity.

### The bacterial and archaeal communities under the canopy of focal tree species exhibited different co-occurrence patterns with increasing tree species richness

The shifts of topological characteristics in co-occurrence network inferred from soil bacteria along a tree species richness gradient suggests that tree species richness influences its complexity, however, contrary to our hypothesis (H2), the network complexity decreased progressively from monocultures to 2-tree species mixtures and 4-tree species mixtures. Bacterial network assembly has been found in many studies to be a deterministic process involving competitive interactions, non-overlapping niches, and thus follows a power-law distribution pattern when bacterial communities are constructed [42–43, 87]. Therefore, we proposed the niche differentiation caused by the tree species richness could be the main reason for changes in bacterial network complexity, with plant monocultures providing a smaller variety of resources and weaker niche differentiation than polycultures, and the weaker niche differentiation, t- he stronger the microbial interactions would be [43, 87–88].

The topological features of archaeal co-occurrence network are not influenced by tree species richness, contrary to our expectations (H2). One proposed explanation is that the archaeal interaction is structured as a random network following the Erdos-Renyi model [43, 89], where the presence or absence of edges is a stochastic process, implying that all interactions between archaea are equally possible. This view is also supported in a recent study of archaeal biogeography showing that the diversity patterns of soil archaea are largely influenced by stochastic processes [90], that is, neutral processes are more important than deterministic factors for soil archaea.

### The roles of bacterial and archaeal community in regulating the relationship between tree-shrub species richness and community-level tree productivity (BEF)

Both tree and shrub species richness contributed significantly to the increase in stand-level tree productivity, confirming our hypothesis (H3). Tree species richness can promote their productivity and thus accelerate C stock [91], and the underlying mechanism is often summarized as ecological niche complementarity [12]. Although shrub competition exists at low shrub species richness levels, but generally, diverse shrub communities positively contribute to stand-level tree productivity, suggesting that competition between shrubs and trees is reduced at higher shrub diversity, and indicating that complementarity effects extend from tree-tree interactions to tree-shrub interactions [16].

In addition, our study also provides insight into the potential role of microbial communities play in this positive BEF relationship. The SEM model suggests that soil C/N is a critical linkage between plant diversity and tree productivity by influencing the bacterial and archaeal communities. Bacteria and archaea inhabiting forest soil are important players in geochemical cycles and organic matter recycling, particularly in the C cycle [92]. The complexity of C cycling is often interlinked with the N cycle, influencing nitrification and denitrification processes and subsequently C/N [93]. Plant species richness significantly drove incremental soil C/N, which can be explained by increased carbon release from trees to the soil through litter production [94] and root exudates [95]. Both C and N are closely linked to the microbial growth and development in biogeochemical cycles, and C/N has a direct effect on the relatively microbial biomass C (MBC/C_org_), mainly because soil bacteria and archaea are predominately heterotrophic organisms that generally derive energy from the decomposition and mineralization of organic matter [36]. In a given ecosystem with high nutrient and resource availability, microbial biomass synthesis is prioritized over catabolism [96]. As a result, the stoichiometry (e.g., C/N) of soil organic matter is critical for regulating microbial communities and increasing microbial activity. Such increases can therefore induce biogeographical cycling of nutrients and maintain higher levels of functioning by increasing physiological potential of microorganisms, thus promoting tree volume growth at the community level. This view is supported by a study showing that plant diversity mediates the metabolic activity of soil microbes via higher root inputs and soil N status and C storage, which would be expected to lead to increased microbials activity [21]. However, in this study, the response of soil bacterial and archaeal communities was only investigated using amplicon sequencing. The response of microbial functions to increased plant species richness would be another intriguing exploration for future research.

## Conclusions

Here, we provided pioneering evidence for the interactive effects of tree and shrub species richness on soil bacterial and archaeal communities under the canopies of focal trees in our long-term biodiversity forest experiments. We demonstrated that *alpha* diversity, co-occurrence networks and community composition of bacteria and archaea follow different patterns towards increasing tree and shrub species richness.

For bacterial communities, the *alpha* diversity and the complexity of co-occurrence network decreases with increasing tree species richness, and the effect of shrub species richness on bacterial *alpha* diversity varied across tree species richness levels. Our results highlighted the dilution effect of tree species richness on soil bacterial diversity in tree diversity experiment. We also demonstrated that changes in bacterial community composition were the result of the direct effects of plant species richness, or indirect effects of them via changing edaphic properties (e.g., C/N and pH). In contrast, for archaeal communities, the effects of tree and shrub species richness on *alpha* diversity, microbial network complexity and community composition were somehow ambiguous, while edaphic properties barely altered the archaeal community composition. Finally, we found that both tree and shrub species richness strongly increased the stand-level tree productivity through direct or indirect regulations on soil microbiota, however, their contributions and the roles of bacterial and archaeal communities in this process were content dependent. Tree species richness could indirectly accelerate bacterial diversity and modulate bacterial community composition via stimulating soil C/N, inducing a cascading effect on the tree productivity. As for archaea, only the diversity of them increased with increasing soil C/N that may be attributable to tree species richness, and thus contributed to stand-level tree productivity. Our findings highlighted the important role of soil microbiome in modulating the relationship between tree and shrub species richness and productivity in subtropical forests.

## List of abbreviations

BEF: Bbiodiversity-Ecosystem Functioning
MBC: Microbial Biomass C
MBN: Microbial Biomass N
QIIME2: Quantitative Insights into Microbial Ecology 2
ASVs: Amplicon Sequence Variants
SEM: Structural Equation Modelling
RMSEA: Root Mean Square Error of Approximation
CFI: Comparative Fit Index
SRMR: Standardized Root Mean squared Residuals
Corg: soil organic C
Norg: soil organic N
SM: soil moisture

## Declarations

### Ethics approval and consent to participate

Not applicable.

### Consent for publication

Not applicable.

### Availability of data and material

Data of seedling biomass, soil physicochemical parameters, and fungal community composition acquired in the study are all included in the manuscript and supplementary material. We submitted the representative sequences from Illumina MiSeq sequencing to NCBI Sequence Read Archive (SRA) database with the accession code PRJNA816566.

## Authors’ contributions

NZ and LQ planned and designed the research. TY collected the soil samples in BEF-China platform and soil physicochemical properties data. S.T. performed data analyses and wrote the draft of the manuscript in close consultation with NZ. GFV, NZ, ST and LQ contributed substantially to the revision of the manuscript. All authors read and approved the final manuscript.

## Acknowledgements

Not applicable.

**Additional file Figure S1.** Flow chart of sampling, DNA extraction, microbial sequencing and the detailed information of dissecting microbial community driven BEF relationships in a subtropical forest.

**Additional file Figure S2.** *A priori* structural equation modeling (SEM) hypothesized causal pathways of how tree/shrub species richness and soil properties may influence stand-level tree productivity through modifying the bacterial and archaeal communities and microbial biomass carbon or nitrogen content.

**Additional file Figure S3.** Soil bacterial and archaeal *alpha* diversity under three tree species richness levels (1, 2, and 4) and four shrub species richness (0, 2, 4, and 8), respectively for both ectomycorrhizal fungi-colonized trees (EcM) and arbuscular fungi-colonized trees (AM).

**Additional file Figure S4.** Principal coordinates analysis (PCoA) with unweighted unifrac distances matrices to visualize the bacterial and archaeal community composition for ectomycorrhizal fungi-colonized trees (EcM) and arbuscular fungi-colonized trees (AM). a. the effect of mycorrhizal types on community compositions of bacteria and archaea. b. the combined effects of tree species richness and shrub species richness on community compositions of bacteria and archaea, respectively for ectomycorrhizal fungi-colonized trees (EcM) and arbuscular fungi-colonized trees (AM).

**Additional file Figure S5.** Pairwise correlation matrix of environmental factors with Mantel tests of bacterial and archaeal communities. Red and blue lines indicate positive and negative correlations, respectively, while solid and dashed lines indicate the significant correlations (*p* < 0.05) and insignificant correlations (*p* > 0.05).

**Additional file Figure S6.** Taxonomic classifications of soil bacterial and archaeal community for ectomycorrhizal fungi-colonized trees (EcM) and arbuscular fungi-colonized trees (AM).

**Additional file Figure S7.** The co-occurrence networks of bacterial communities in three tree species richness levels (1, 2, and 4) coupled with four shrub species richness levels (0, 2, 4, and 8). The nodes in the networks are colored referred to the taxonomic assignments at phylum level and the size of each node is proportional to the relative abundance.

**Additional file Figure S8.** The co-occurrence networks of archaeal communities in three tree richness levels (1, 2, and 4) coupled with four shrub richness levels (0, 2, 4, and 8). The nodes in the networks are colored by the taxonomic assignments at phylum level and the size of each node is proportional to the relative abundance.

**Additional file Table S1.** The overview of sample information across the experimental plots.

**Additional file Table S2.** The linear mixed effect model summaries for main effects of tree and shrub species richness on the community structure of bacteria and archaea, respectively for ectomycorrhizal fungi-colonized trees (EcM) and arbuscular fungi-colonized trees (AM).

**Additional file Table S3.** The significantly differential abundances of bacterial and archaeal phyla under the combined effects of tree and shrub species richness by Generalized Linear Model tests.

**Additional file Table S4.** The significantly differential abundances of bacterial and archaeal amplicon sequence variants (ASVs) assigned to functional groups under the combined effects of tree and shrub species richness by Generalized Linear Model tests.

