## Supplementary figures and images for "Tree and shrub richness modify subtropical tree productivity by modulating the diversity and community composition of soil bacteria and archaea"

### Additional file Figure S1

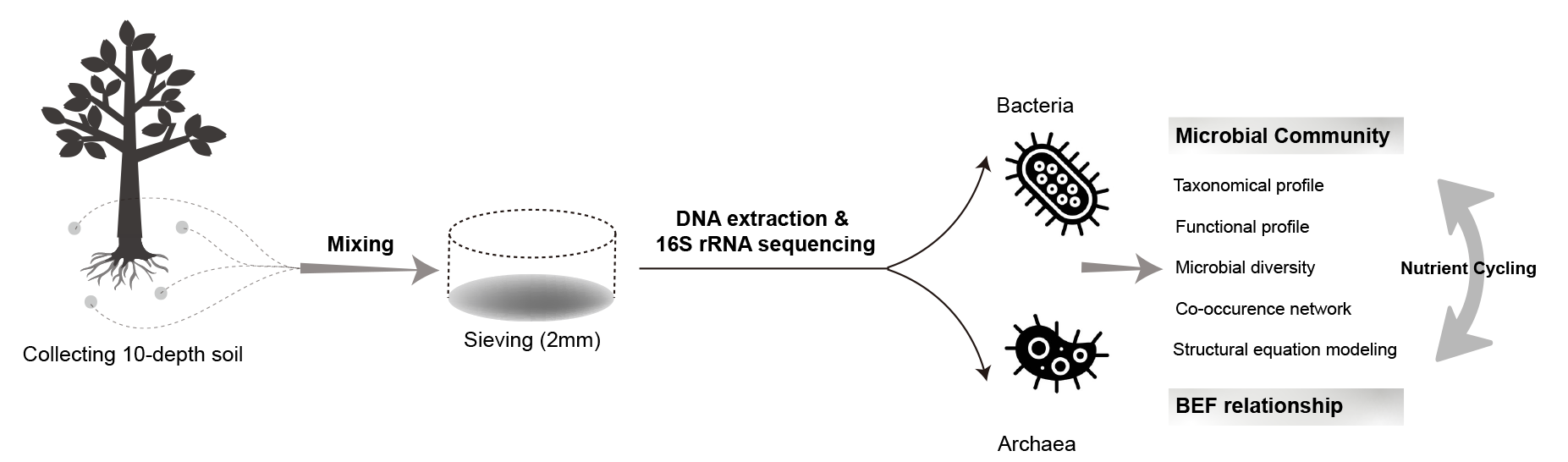

### Additional file Figure S2

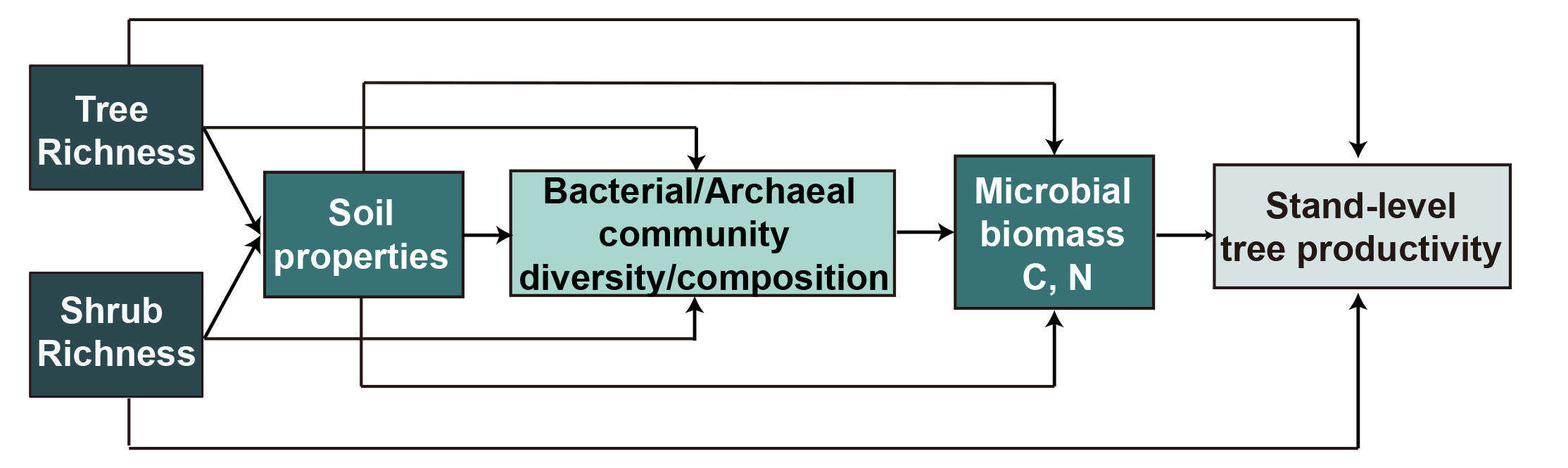

### Additional file Figure S3

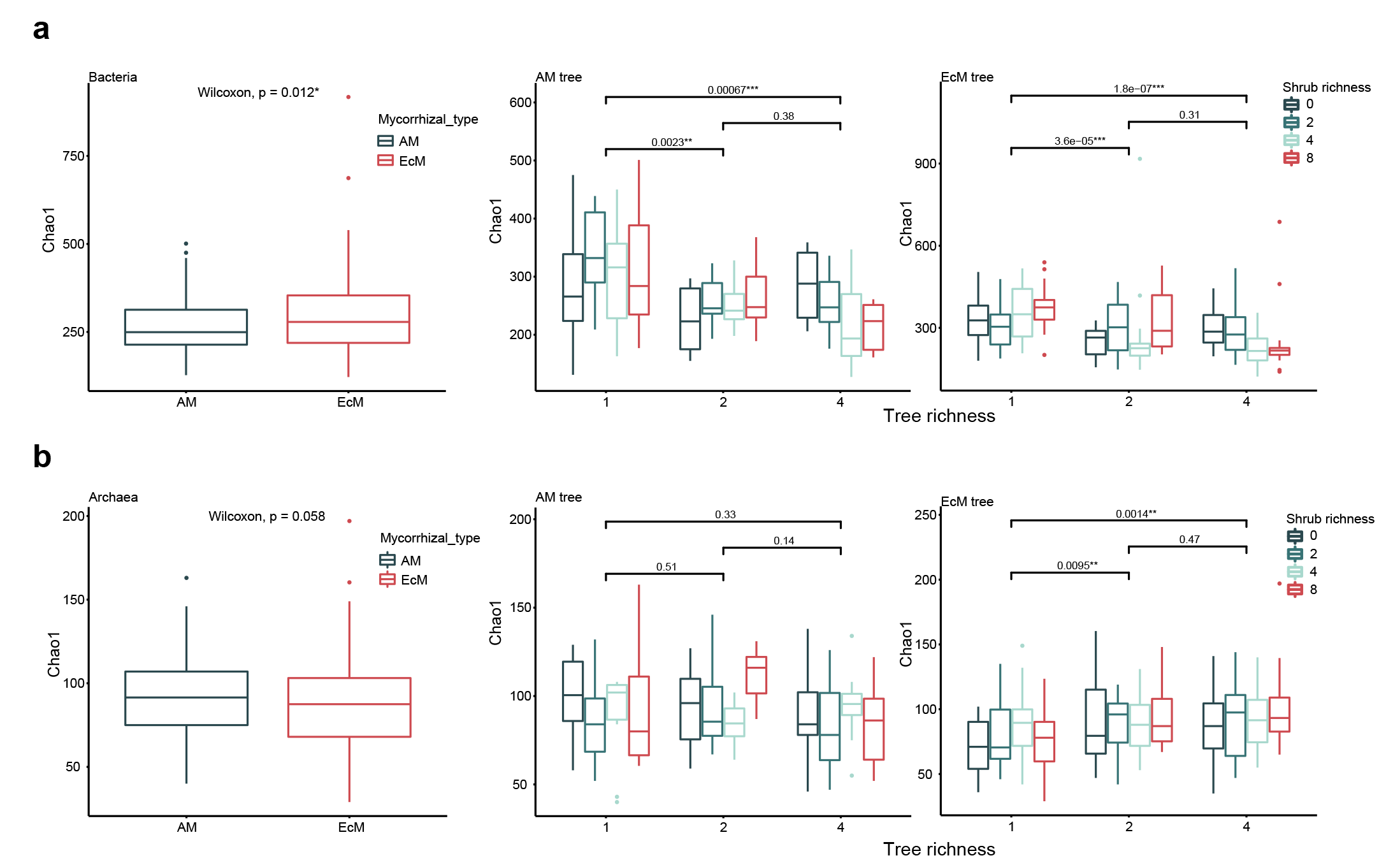

### Additional file Figure S4

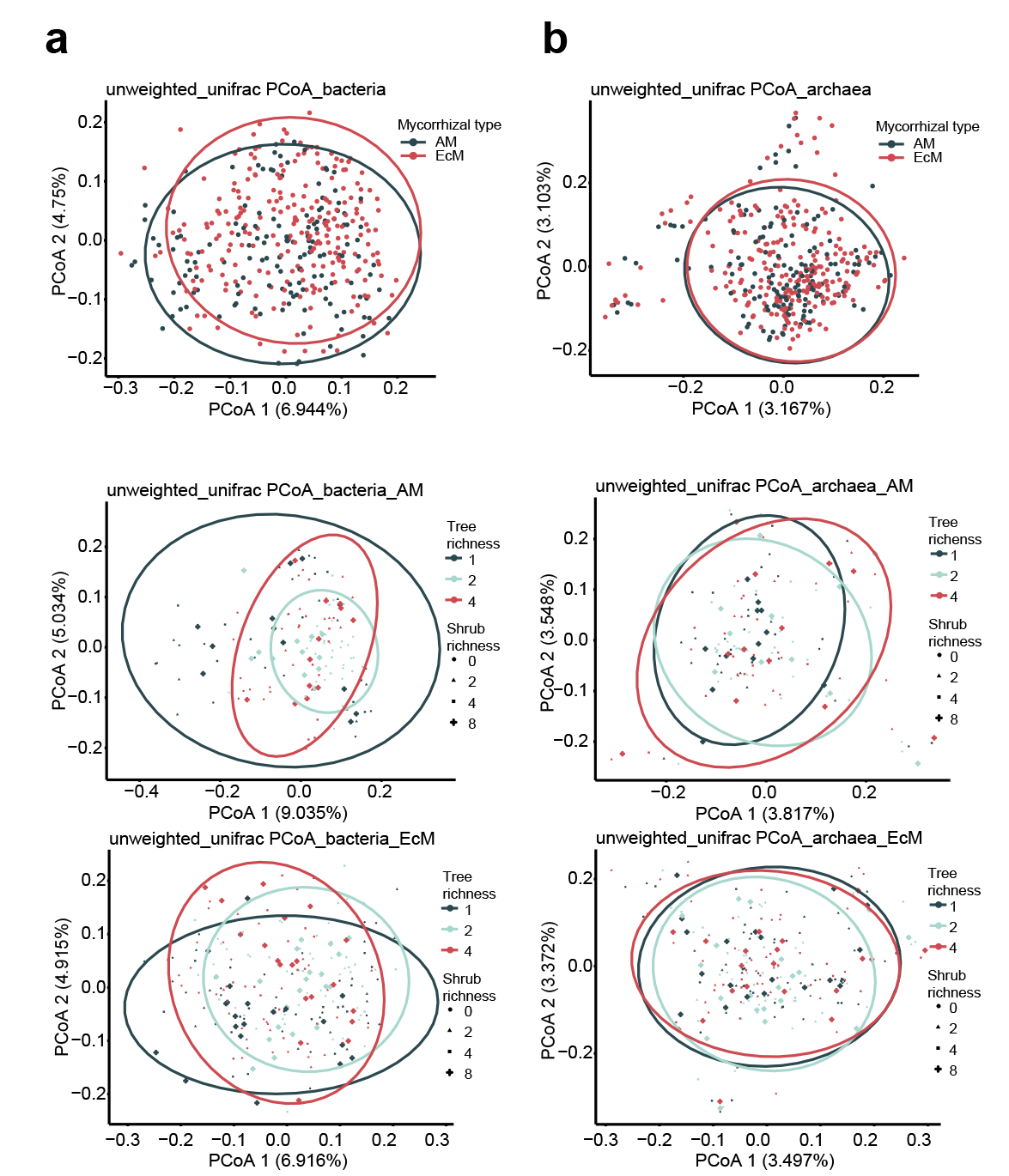

### Additional file Figure S5

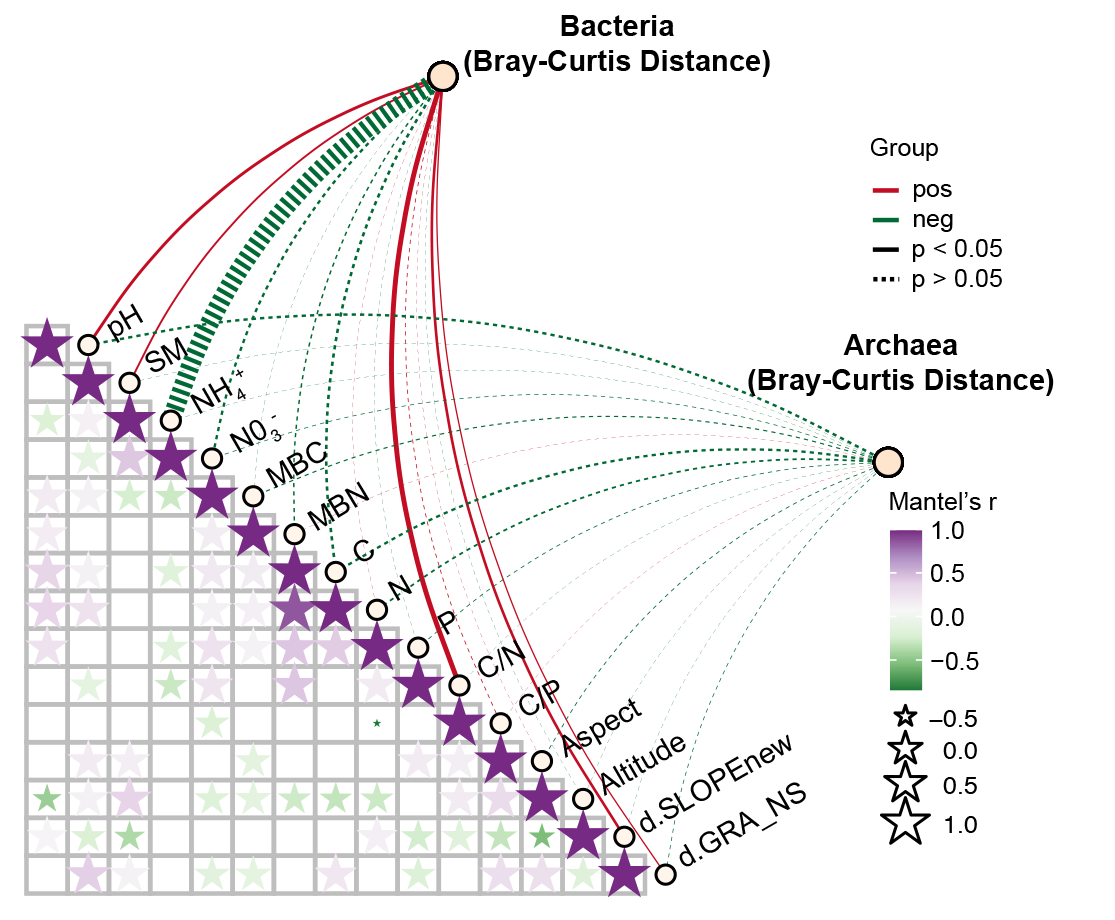

### Additional file Figure S6

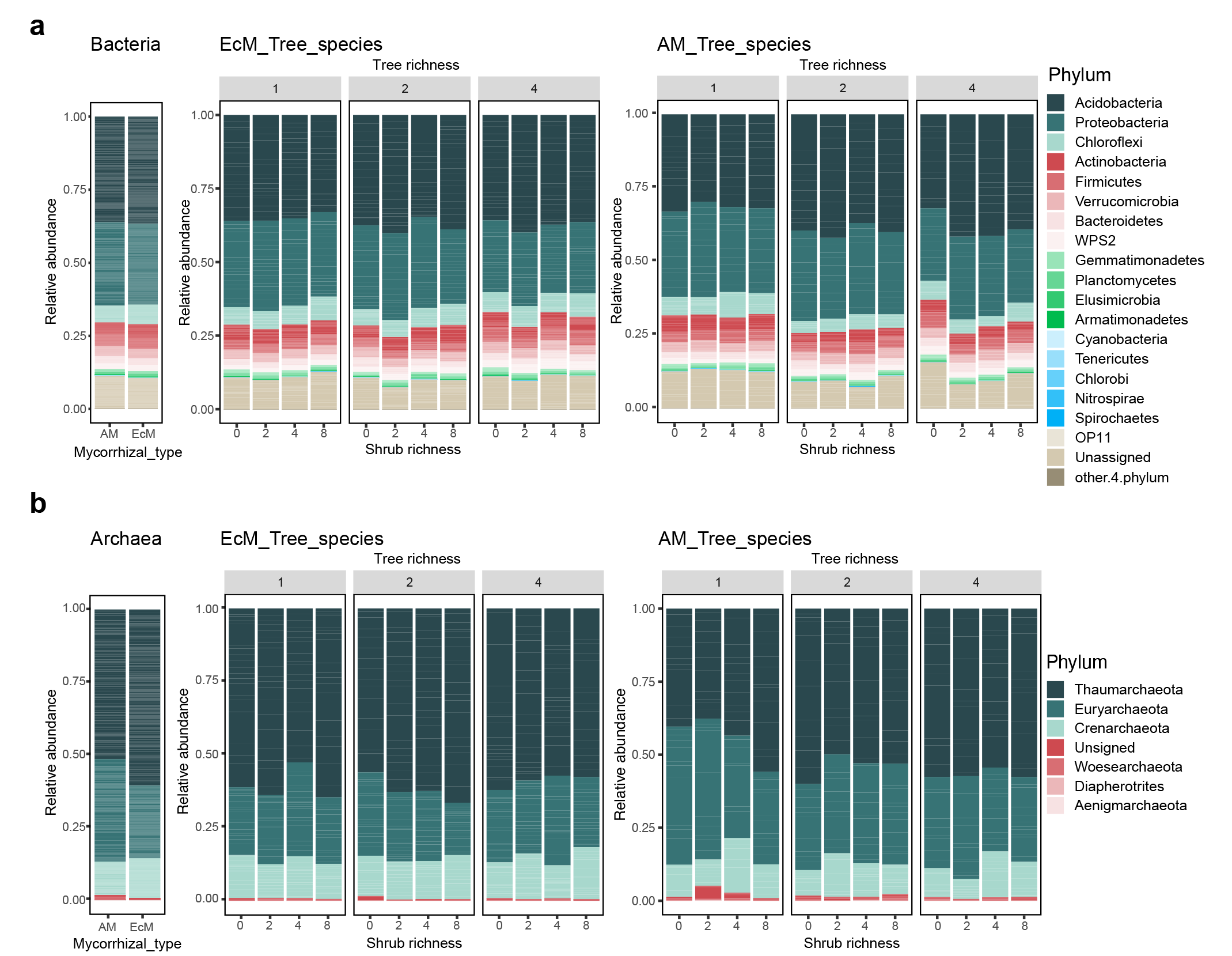

### Additional file Figure S7

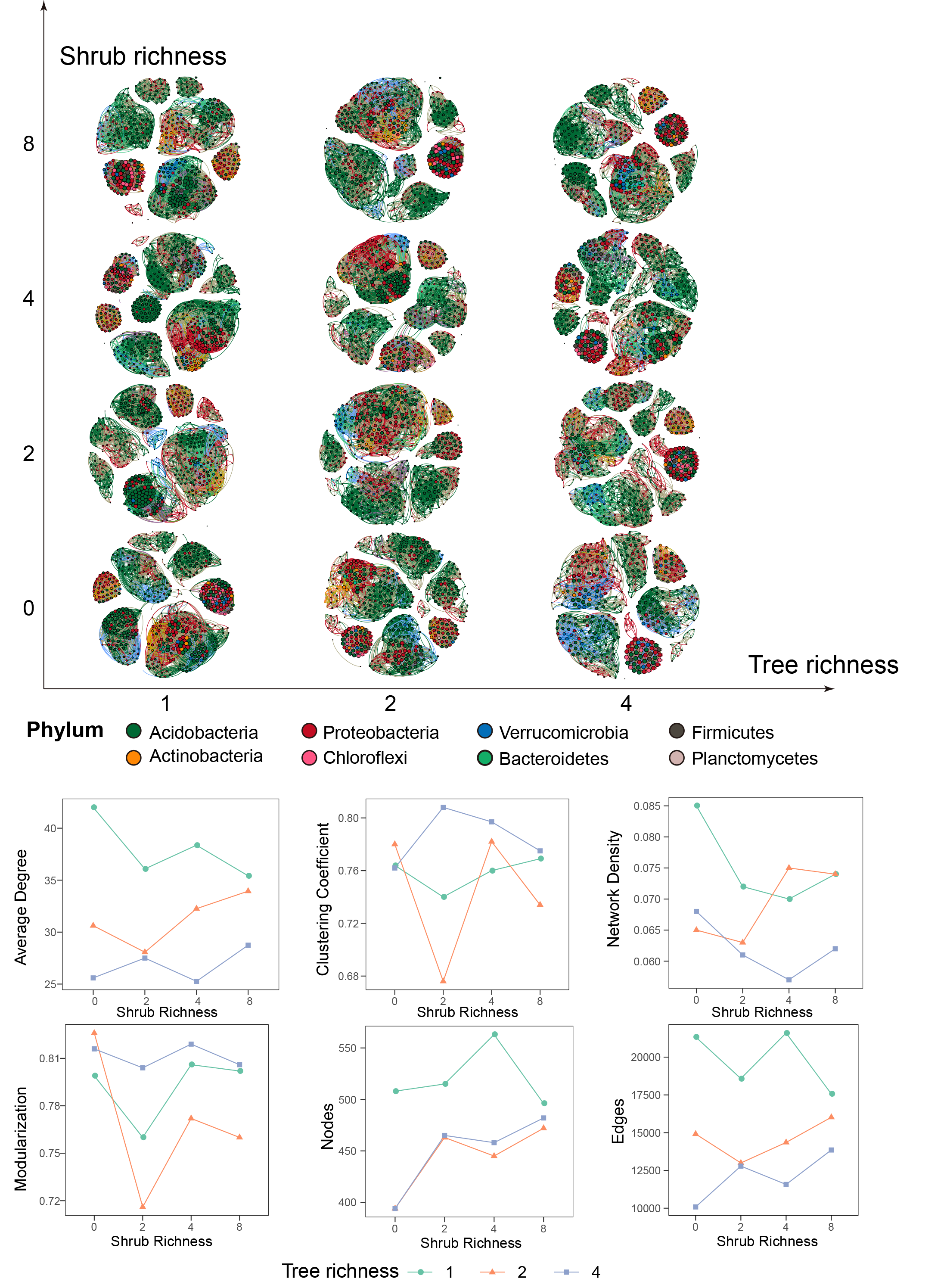

### Additional file Figure S8

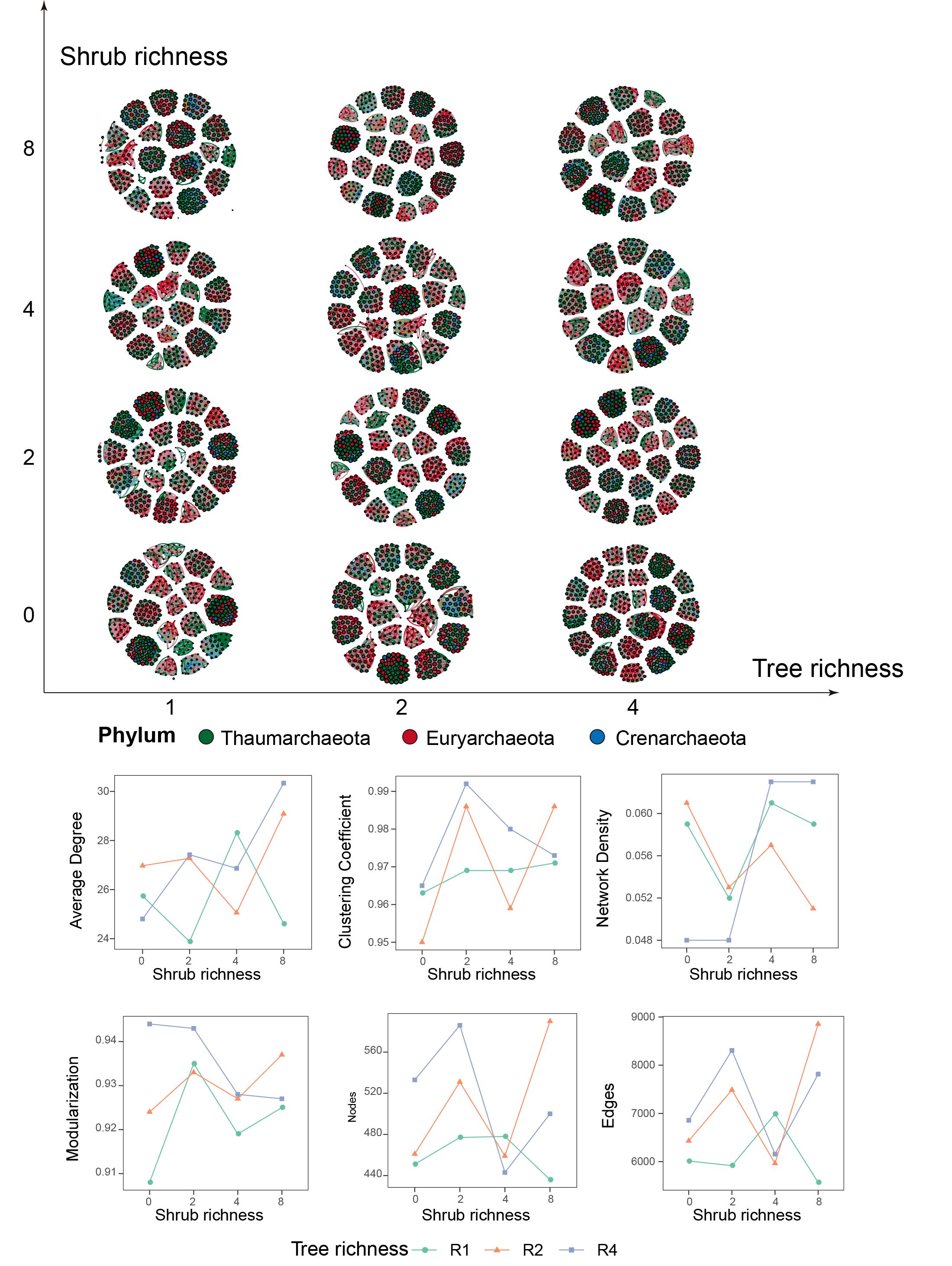
